# Mdm2 restrains the activity of the CRL4^Cdt2^ E3 ubiquitin ligase to promote cell cycle progression through the G2/M phase

**DOI:** 10.1101/2025.07.09.663887

**Authors:** Humaira Shah, Auqib Manzoor, Tabasum Ashraf, Ashraf Dar

## Abstract

The canonical function of the Mdm2 oncoprotein is widely recognized as the negative regulation of the p53 tumor suppressor. However, growing evidence indicates that its physiological activities extend far beyond p53. Here, we show that Mdm2 promotes cell cycle progression at the G2/M phase through the ubiquitin-mediated degradation of the substrate recognition adaptor Cdt2 of the CRL4^Cdt2^ E3 ubiquitin ligase complex, independently of p53. The attenuation of CRL4^Cdt2^ activity by Mdm2 stabilizes its cell cycle-specific substrates including p21, Set8, and Cdt1, at the G2/M phase following their proteasomal degradation in the S phase. Furthermore, the delay in cell cycle progression at the G2/M phase and the decreased cell proliferation observed in the absence of Mdm2 are largely caused by an increase in Cdt2 and a subsequent decrease in p21. Collectively, our data illustrate a previously unexplored mechanism by which Mdm2 regulates the cell cycle and promotes cellular proliferation by neutralising the CR4^Cdt2^ E3 ubiquitin ligase activity and subsequently stabilizing p21.

## Introduction

Mdm2 is an oncoprotein whose oncogenic potential primarily stems from its ability to ubiquitinate and degrade the tumor suppressor p53 (Haupt et al. 1997). Under normal, unstressed conditions, Mdm2 maintains cellular homeostasis by keeping p53 levels low. However, in response to genotoxic or oncogenic stress, the Mdm2-p53 interaction is disrupted, leading to p53 stabilization and activation of downstream targets, including the cyclin-dependent kinase inhibitor p21 and pro-apoptotic genes such as *BAX*, *PUMA*, and *NOXA* (Vousden and Prives 2009)(Riley et al. 2008). Depending on the nature and severity of the damage, this activation can trigger cell cycle arrest, senescence, or apoptosis (Vousden and Lane 2007).

While the canonical Mdm2-p53 axis is well characterized in DNA damage responses, emerging evidence suggests that Mdm2 also carries out critical p53-independent functions, particularly under physiological, non-stressed conditions. For example, Mdm2 has been implicated in promoting DNA replication via degradation of PARP1 (Giansanti et al. 2022), supporting cancer cell growth and survival through targeting of p73 (Stommel and Wahl 2004)(Klein et al. 2021a), and paradoxically inhibiting cell cycle progression via degradation of Cdc25C (Stark and Taylor 2006)(Giono et al. 2017). These findings broaden our understanding of Mdm2’s physiological roles beyond p53 regulation.

Central to cell cycle arrest following Mdm2-p53 disruption is the transcriptional activation of p21, which enforces G1/S phase arrest by binding and inhibiting Cdk2, thereby blocking DNA synthesis (El-Deiry et al. 1993)(Wade Harper 1993). Interestingly, under basal conditions, p21 expression can also occur independently of p53, highlighting the role of alternative regulatory mechanisms (Abbas and Dutta 2009)(Gartel and Radhakrishnan 2005). Among these, post-translational regulation via the CRL4^Cdt2^ E3 ubiquitin ligase complex plays a key role. CRL4^Cdt2^ targets p21, along with Set8 and Cdt1, for proteasomal degradation at the G1/S transition (Abbas et al. 2008)(Centore et al. 2010)(Jin et al. 2006)(Kim et al. 2008). Timely degradation of these substrates is critical for S-phase progression and maintenance of genomic integrity (Lovejoy et al. 2006)(Abbas and Dutta 2011). Notably, p21, Set8, and Cdt1 reaccumulate during G2/M (Coleman et al. 2015), suggesting that suppression of CRL4^Cdt2^ activity is required for mitotic entry. However, the mechanisms underlying this suppression in actively cycling cells is not known.

In this study, we uncover a novel, p53-independent Mdm2-CRL4^Cdt2^-p21 axis that governs the G2/M transition under unstressed conditions. Using CRISPR-Cas9- mediated knockout of Mdm2 in human osteosarcoma U2OS cells, we show that loss of Mdm2 leads to a pronounced G2/M arrest and decreased cellular proliferation, phenotypes that persist despite forced p53 depletion. We further demonstrate that Mdm2 attenuates CRL4^Cdt2^ activity by ubiquitin-mediated degradation of Cdt2, allowing the stabilization of p21 (and other substrates: Set8 and Cdt1) during G2/M- a critical step for mitotic entry. Notably, ectopic expression of p21 alone is sufficient to rescue the G2/M block in Mdm2-deficient cells, establishing p21 as a central effector of this newly defined regulatory axis. These findings align with recent studies that highlight pro-proliferative and tumor-promoting roles for p21, including its ability to inhibit apoptosis and support mitotic progression (Roninson 2002)(Abbas and Dutta 2009)(Gartel and Tyner 2002)(Blagosklonny 2002). Conclusively, this study provides mechanistic insight that distinguishes Mdm2’s function from the canonical p53 axis, highlighting its critical role in facilitating orderly mitotic entry under non-stressed conditions. These findings underscore the potential of targeting Mdm2 therapeutically, even in p53-deficient cancers (Wade et al. 2013)(Burgess et al. 2016).

## Materials and Methods

### Cell culture and transfections

Cells were maintained in 6-well plates, 60mm or 100mm dishes. HEK 293T, U2OS, and HeLa and grown in Dulbecco’s Modified Eagle Medium (DMEM, Gibco-Life Technologies) with 10% Fetal Bovine Serum (FBS; Gibco-Life Technologies) and 1% penicillin-streptomycin. HCT-116 Cells were grown in McCoy’s 5a medium (Gibco-Life Technologies) supplemented with 10% FBS. Cells were split by trypsinization using 0.25% Trypsin-EDTA (Gibco-Life Technologies). Plasmids were transfected into cells at 60% confluency using polyethyleneimine reagent (Polyscience) as suggested by the manufacturer.

### Generation of Mdm2 KO cell line

Small guide RNAs(sgRNA) were selected to generate Mdm2 knockout U2OS cells following careful analysis of the transcript using both NCBI and Ensemble and identified using www.benchling.com/crispr. Optimal sgRNA pairs were identified with a low combined off-targeting score (Table SI) . dsDNA sg oligos were ligated into BbsI-digested target vectors PX458 and PX459. U2OS cells at 60% confluency were co-transfected with pSpCas9(BB)-2A-GFP (PX458) and pSpCas9(BB)-2A-Puro (PX459) carrying sg Mdm2 using PEI according to the manufacturer’s instructions. 24 hours after transfection, the medium was replaced with a fresh medium containing 5µg/ml of puromycin. After 48 hours of puromycin selection, the medium was replaced again with a fresh medium without puromycin to recover for 48 h. Single cells were placed in individual wells of a 96-well plate and grown up to ∼80% confluency, and then transferred into six-well plates. After reaching ∼80% confluency, the clones were screened for MDM2 knockout by T7 endonuclease assay, and expression was analysed by immunoblotting.

### shRNA plasmid constructs

Short hairpin RNA (shRNA) sequences targeting human MDM2, Cdt2, p53, p21, set8, and Cdt1 were designed using the http://www.oligoengine.com. Oligonucleotides encoding the shRNA were purchased from Sigma-Aldrich. The sense and antisense strands were annealed and ligated into the pSUPER vector at the restriction enzyme sites HindIII and BglII, downstream of the H1-RNA promoter. Plasmid constructs were confirmed by restriction digestion, and knockdown efficiency was checked by western blotting. The oligo sequences for different shRNA are given in Table S1.

### Western blotting and antibodies

For western blotting, cells were lysed with the requisite amounts of radioimmunoprecipitation assay (RIPA) buffer containing 1X protease inhibitor cocktail (Sigma) and centrifuged at 13,000 g at 4°C for 15 min. Then, 2X sample buffer was added to the protein supernatants, and the samples were boiled at 100°C for 5 min. Protein samples were resolved under denaturing conditions on SDS-PAGE and transferred onto nitrocellulose membranes (Sigma). Membranes were blocked with 5% non-fat dried milk, incubated overnight at 4°C with primary antibodies, followed by washes and incubation with HRP-conjugated secondary antibodies (Dar et al. 2013). Immunoreactive bands were visualized using chemiluminescence reagents luminol and coumaric acid (Sigma) and detected with a Chemi Doc XP imaging system (Bio-Rad).

The following antibodies were used: β-actin (Santa Cruz Biotechnology; Sc-58673; 1:1000), Cdt2 (Bethyl Laboratories; A301-109A; 1:500), FLAG (Cell signaling technology; #2368;1:1000) Myc (Cell signaling technology; #2276;1:1000), Cdt1(Cell signaling technology;#8064; 1:500), p21(Cell signaling technology; #2947; 1:500), set8 (Cell signaling technology; #2996;1:500), p53(Santa Cruz Biotechnology; 1:1000; sc-126), Cul4A(Cell signaling technology; #2699; 1:500), Mdm2(Cell signaling technology; #86934;1:500), DDB1(Cell signaling technology; #5428;1:500 CST), Cyclin B1(Cell signaling technology; #12231;1:500), Cyclin A2(Cell signaling technology; #4656;1:500), Cyclin D1(Cell signaling technology; #55506;1:500), HA (Cell signaling technology; #23676;1:500)

### Coimmunoprecipitation

Cells were lysed on ice for 30 min in a lysis buffer (50 mM Tris-Cl [pH 8.0], 10% glycerol, 150 mM NaCl, 1 mM EDTA, 0.1% Triton X-100, 1 mM dithiothreitol (DTT), 50 mM NaF, 1 mM Na_3_VO_4_, 1 mM glycol phosphate, 1X protease inhibitor cocktail), followed by centrifugation at 15,000 rpm for 10 min. Sepharose G beads(Invitrogen) were mixed with the requisite antibody and incubated at 4^°^C overnight. Cell extracts were mixed with pre-treated beads and kept for mixing at 4^°^C for 4-6 hours. The beads were washed with ice-cold 1X PBS three times and a final wash with lysis buffer. 2X SDS loading buffer was then added to the beads, and samples were boiled at 100^°^C for 5 minutes for western blot analysis (Dar et al. 2014).

### Ubiquitination assay

For the *in vivo* ubiquitination assay, the plasmid-transfected 293T cells were treated with MG132 (40 μM) for 1 h before harvesting. Cells were harvested in denaturing ubiquitination buffer (50 mM Tris-Cl [pH 8.0], 5 mM DTT, and 1% SDS) and immediately boiled for 10 min at 95°C, followed by cooling on ice for 10 min. The cell lysates were sonicated, and the supernatant was recovered after centrifugation at 15,000 rpm for 20 min. The supernatant was diluted with 9 volumes of buffer containing 50 mM Tris-HCl (pH 8.0), 150 mM KCl, 5% glycerol, 0.4% NP-40, and protease inhibitors and subjected to immunoprecipitation followed by western blotting (Shibata et al. 2014).

### Half-life measurements

For the cycloheximide chase assay, cells were treated with freshly prepared cycloheximide (5 mg/ml in H_2_O) and harvested at different time points, followed by western blotting of the cell lysates (Kiran et al. 2018).

### Real-time q-PCR

Total RNA was isolated from cells by using TRIzol reagent (Invitrogen), following the manufacturer’s instructions, and was used for cDNA synthesis with cDNA synthesis kit (Invitrogen). The cDNAs were used as the templates for real-time quantitative PCR (q-PCR) using SYBR green PCR master mix (Invitrogen) and Bio-Rad CFX Opus 96 Real-Time PCR System. The expression of *GAPDH* was used as the internal control.

Bar graphs represent the relative ratios of target genes to *GAPDH* gene values. For each biological sample, triplicate reactions were analysed. The sequences of primers used for RT-PCR analysis are given (5′→3′) in the supplementary Table S1.

### Cell cycle analysis

Cells were grown to a 50-60% confluence, harvested by trypsinization, and washed with 1X PBS. The cells were removed by vigorous pipetting into 15 ml tubes and pelleted at 1000 rpm for 5 minutes. For fixing, the pellets were washed once with 1x PBS supplemented with 1% serum and centrifuged again at 1000 rpm for 5 minutes. The pellet was resuspended in 500μl of 1X PBS and fixed by the addition of 5 mL of ethanol drop-wise while vortexing to ensure the disaggregation of cell clumps into a single cell suspension. Finally, the cells were stored at 4°C for at least overnight in the dark. For cell cycle analysis in flow cytometer, the cells were centrifuged at 1000 rpm for 5 minutes, washed once with PBS (+ 1% serum), spun again at 1000 rpm for 5 minutes and then resuspended in 200-500μl of propidium iodide/ RNase solution (50μg/ml propidium iodide, 10mM Tris, pH7.5, 5mM MgCl2 and 20mg/ml RNase A) and incubated at 37°C for 30 minutes. Samples were then subjected to flow cytometry using BD FACS Aria, and statistical analysis was done using FlowJo v10.

### Cell Viability

After growing different cells for different experiments, the cell viability was measured by MTT assay. 5mg/ml MTT(Sigma-Aldrich) was added to each well for 4 h at 37°C. After incubation, 1 mL of DMSO (solubilizing reagent) was added to each well, agitated at a low speed for 10 min. The absorbance value of each well was measured at 570 nm using NanoDrop 1000 Spectrophotometer (Thermo Scientific). Experiments were conducted in triplicate.

## Results

### Mdm2 regulates the G2/M phase of cell cycle independently of p53

The embryonic lethality in mouse caused by deleting Mdm2 gene can be rescued by codeletion of p53 (De Oca Luna et al. 1995)(Jones et al. 1995). However, the requirement of Mdm2 for p53 null cells for survival and proliferation indicates that Mdm2 may be important for cell proliferation independently of p53 (Feeley et al. 2017).

To gain insight into the role of Mdm2 in cell cycle progression, we used CRISPR-Cas9 technology to genetically knock out the Mdm2 gene in U2OS cells. The KO cell line established from single cell derived clone was chosen for further investigation (**Fig 1A**). As expected, genetic ablation of Mdm2 led to a concomitant stabilization of p53 (**Fig. 1A, S1A–B**). Given that upregulated p53 promotes cell cycle arrest and impairs cellular proliferation through the transcriptional activation of *p21* under DNA damage or stress conditions, we sought to assess how Mdm2 knockout (KO) cells with elevated p53 behave during the cell cycle under normal (unstressed/undamaged) conditions.

**Fig. 1.**
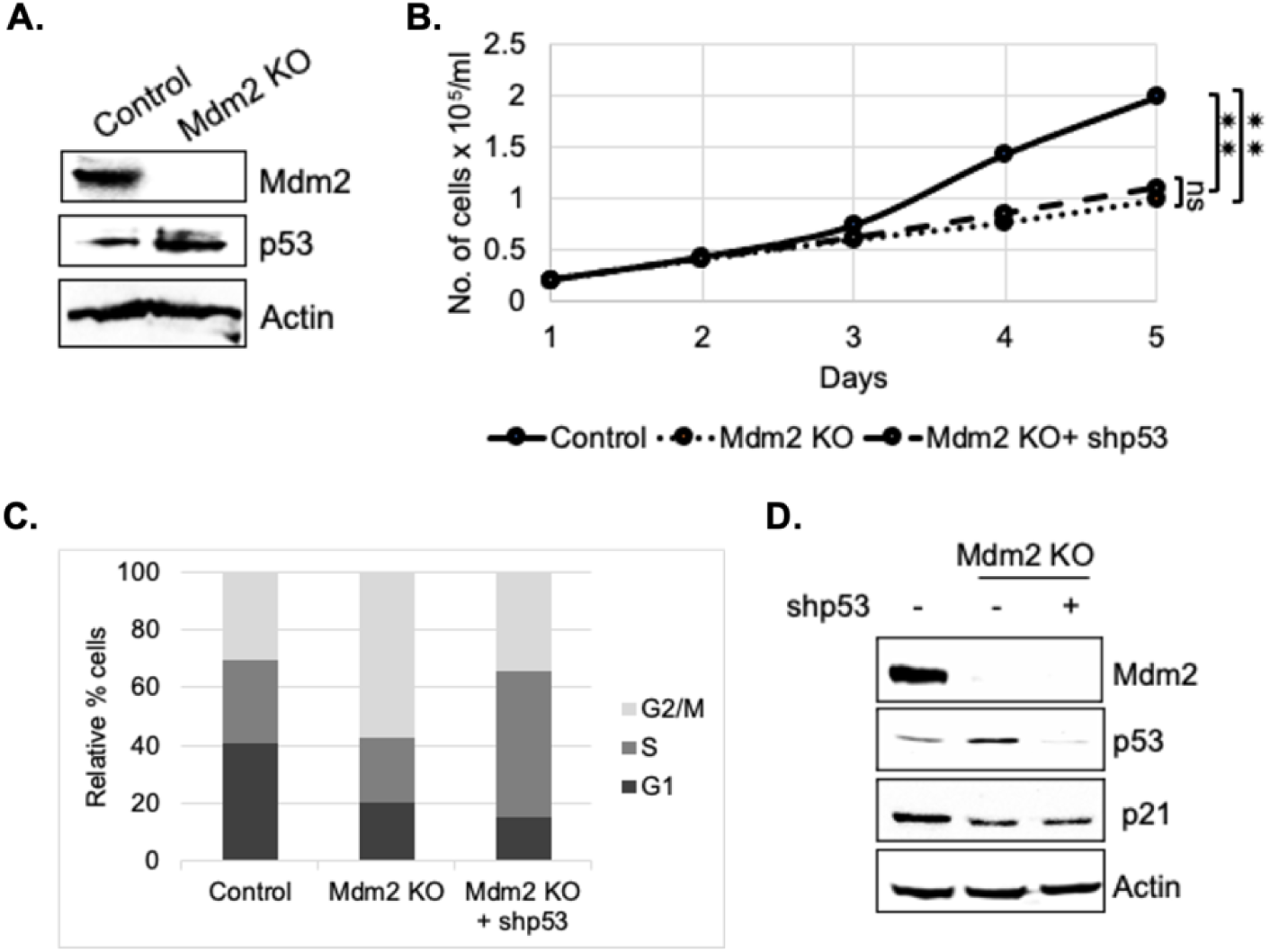
Mdm2 is essential for viability of cells independently of p53. **(A)** Mdm2 was Knocked out in U2OS cells using CRISPR-Cas9. The knockout clone was confirmed with downregulation of Mmd2 and a concurrent upregulation of p53 using western blotting of lysates using indicated antibodies. **(B)** Proliferation rates of wild type, Mdm2 KO and Mdm2 KO+ shp53 U2OS cells. Mean ± SD of 3 replicates. *, *P* < 0.01, two- tailed Student’s *t* test. (**C**) FACS analysis of different cell cycle phases in control, Mdm2 KO and Mmd2 KO+shp53 U2OS cells. (**D**) U2OS cell lysates, from experiments performed in parallel to those whose results are shown in panel C, were probed with the indicated antibodies.

Deletion of Mdm2 gradually slowed cellular proliferation (**Fig.1B**). FACS analysis of Mdm2-deficient cells revealed an increased G2/M population and a significant reduction in G1-phase cells (**Fig. 1C**). To determine whether the G2/M arrest and reduced proliferation were due to the absence of Mdm2 itself or the concurrent increase in p53 levels, we silenced p53 using shRNA and monitored cell proliferation and cell cycle progression. Notably, depletion of p53 in the Mdm2 KO background did not rescue the G2/M arrest (**Fig. 1C**) or restore normal proliferation significantly (Fig 1B). Instead, the depletion of p53 in Mdm2 KO background arrested the cells in S-phase.

Since *CDKN1A*, which encodes p21, is a well-known transcriptional target of p53 (Sullivan KD, 2017) and a mediator of its cell cycle inhibitory effects under stress conditions, we examined p21 levels in Mdm2 knockout (KO) cells with elevated p53 to determine whether the reduced cell proliferation observed in these cells was due to elevation in p21 levels. Surprisingly, western blot analysis revealed that p21 levels were downregulated in Mdm2 KO cells despite elevated p53 levels, compared to wild-type cells (**Fig. 1D**). Furthermore, p53 knockdown did not further reduce p21 levels in the Mdm2 KO background (**Fig. 1D**).These findings suggest that Mdm2 is required for p21 stability in a p53-independent manner under unstressed conditions.

Taken together, our results indicate that Mdm2 is essential for cell proliferation and G2/M phase progression independently of p53, possibly by regulating an alternative pathway involved in p21 turnover during the cell cycle.

### Mdm2 regulates master regulator CRL4^Cdt2^ of cell cycle via Cdt2

Previous studies have shown that p21 and other cell cycle regulators, including Cdt1 and Set8, are targeted for degradation by the CRL4^Cdt2^ E3 ligase during S-phase for smooth progression of cell cycle (Havens and Walter 2011). However, these substrates reappear during the G2/M phase (Coleman et al. 2015). Given that Mdm2 is critical for the stability of p21 (**Fig. 1E**), we hypothesised that Mdm2 protein may negatively regulate CRL4^Cdt2^ to stabilize its downstream substrates, including p21, Set8 and Cdt1.

The KO of Mdm2 leads to the accumulation of DDB1 and Cdt2, components of the E3 ligase complex, CRL4^Cdt2^ (**Fig. 2A**). Since both DDB1 and Cdt2 are parts of the same enzyme complex, we focused further investigation on how Mdm2 regulates CRL4^Cdt2^ stability via Cdt2. Furthermore, the stabilization of Cdt2 in response to Mdm2 depletion using specific *sh*RNA in HeLa and HCT116 cells rules out the possibility that the negative regulation of Cdt2 by Mdm2 observed earlier in U2OS Mdm2 KO cells is a cell line-specific effect (**Fig 2B**). Notably, the accumulation of Cdt2 is not due to transcriptional activation, as its its mRNA levels remain largely unchanged following Mdm2 deletion (**Fig. 2C**).

**Fig. 2.**
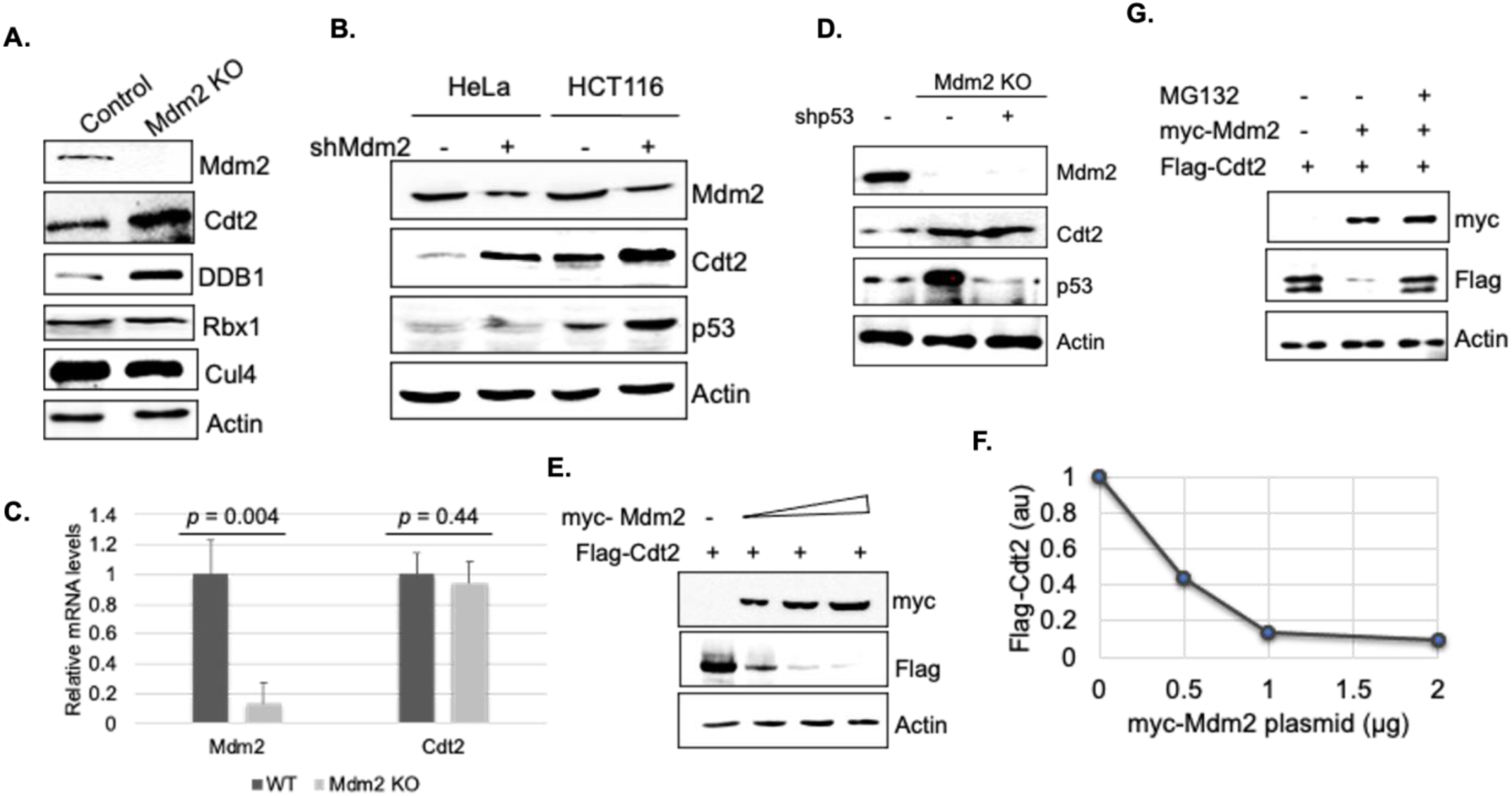
Mdm2 destabilizes CRL4^Cdt2^ E3 ligase. (**A**) Control (wild-type) and Mdm2 KO U2OS cell lysates were immunoblotted as indicated. (**B**) HeLa and HCT116 cells were transfected with control and shRNA against Mdm2 and harvested after 60 h. The cell lysates were probed with antibodies as indicated. (**C**) Mdm2 and Cdt2 mRNA levels in wild type (WT) and Mdm2 KO cells. The transcripts were measured by qRT-PCR and normalized to *GAPDH* mRNA. Mean ± SD of three replicates. *P* values determined are as indicated. (**D**) Western blot analysis of different proteins in lysates prepared from U2OS control cells, Mdm2 KO and Mmd2 KO + shp53 (60h) cells. **(E**) Mdm2 decreases Flag-Cdt2 levels in cells. The results of western blot analysis of 293T cells transfected with plasmids expressing Flag-Cdt2 alone or Flag-Cdt2 plus increasing amounts of myc-Mdm2 are shown. (**F**) The Flag-Cdt2 signal described in the panel E legend was quantitated and plotted relative to the actin signal to measure the destabilization of Flag-Cdt2 with or without coexpressed myc-Mdm2. (**G**) 293T cells overexpressing Flag-Cdt2 and myc-Mmd2 were treated with dimethyl sulfoxide (DMSO) or MG132 for 2 h, as indicated and total cell lysates immunoblotted.

To determine whether concurrent accumulation of p53 upon Mdm2 KO contributes to Cdt2 stabilization, we silenced p53 using shRNA in these cells. The results show that neither stabilized p53 nor its forced depletion affects Cdt2 stability in Mdm2 KO background (**Fig. 2D**), indicating that Mdm2-mediated destabilization of Cdt2 is independent of p53.

Since Mdm2 is known to be amplified in 36% sarcomas and overexpressed in other cancers (Oliner et al. 1992)(Oliner et al. 2016), we examined the effect of Mdm2 overexpression on Cdt2 stability. U2OS cells were transiently transfected with plasmid expressing myc-Mdm2. This overexpression significantly decreased the steady-state levels of Cdt2 (**Fig. 2E-F**). Mdm2-mediated destabilization of Cdt2 was also observed in other cell lines, including 293T, HCT116, and HeLa (**Fig. S2A-C**). Furthermore, this effect is proteasome-dependent, as treatment with the proteasome inhibitor MG132 prevented Cdt2 degradation by transiently overexpressed Mdm2 (**Fig. 2G**).

Overall, these results indicate that Mdm2 regulates the stability of CRL4^Cdt2^ E3 ligase by targeting Cdt2 via proteasome in a p53-independent manner.

### Mdm2 accelerates the degradation of Cdt2 independent of threonine 464 phosphorylation

To assess whether Mdm2-mediated proteasomal degradation of Cdt2 affects protein turnover, we measured the half-life of Cdt2 following translation inhibition using cycloheximide (CHX). Under normal conditions, Cdt2 exhibits a half-life of approximately 50 minutes, which extends beyond 120 minutes upon genetic deletion of Mdm2 (**Fig. 3A-B**). Conversely, ectopic overexpression of Mdm2 accelerates the degradation of both endogenous Cdt2 and exogenous Flag-tagged Cdt2, thereby reducing their half-lives significantly (**Fig. 3FC-D** and **S3A-B**).

**Fig. 3.**
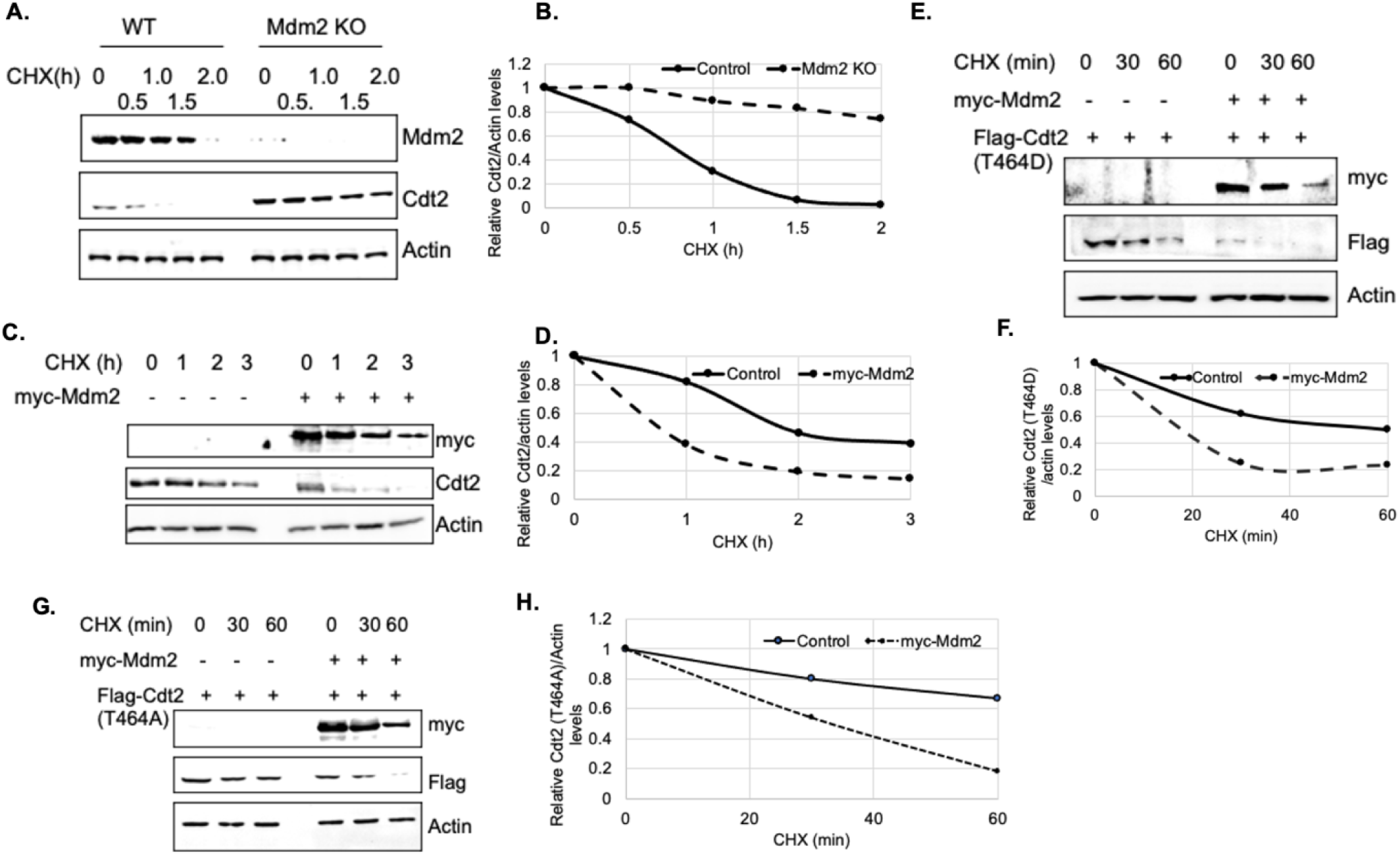
Mdm2 decreases half-life of Cdt2 independent of T464 phosphorylation. (**A**) Mdm2 KO increases the half-life of Cdt2. Wild type (WT) and Mdm2 KO U2OS cells were treated with 100 μg/ml cycloheximide (CHX) for the time points as indicated. The cell lysates were probed with antibodies indicated. (**B**) The Cdt2 bands in panel A were quantitated and normalized to actin, followed by the plotting of results graphically. (**C**) U2OS cells were transfected with empty or myc-Mdm2 plasmid followed by treatment with CHX for different time points. The cell lysates were probed with antibodies indicated. The Cdt2 bands were quantitated using densitometry, normalised to actin and plotted graphically in **D**. (**E**) The plasmid encoding phospho- mimetic (T464D) version of Flag-Cdt2 was coexpressed in 293T cells with empty or myc-Mdm2 plasmids followed by treating cells with CHX for different time points. The cell lysates were probed with antibodies shown. (**F**) The Flag- signals in panel E were quantified, normalized to actin and plotted graphically. (**G**) Flag-Cdt2 (T464A) was overexpressed with empty or myc-Mdm2 plasmids. Cells were treated with CHX for the time points as indicated. The lysates were analysed in western blotting using antibodies indicated. (**H**) The Flag- and actin signals in panel **G** were quantified using densitometry. The values from ratio of Cdt2/Actin were plotted graphically.

Cdt2 is also known to be targeted for degradation by FBXO11 during cell cycle exit, a process inhibited by Cyclin B-Cdk1/Cyclin A-Cdk2-mediated phosphorylation at threonine 464 (T464) (Rossi et al. 2013). To determine whether Mdm2-mediated degradation of Cdt2 is similarly prevented by this phosphorylation, we overexpressed phosphomimetic Flag-Cdt2 (T464D, resistant to FBXO11-mediated degradation) and phosphoresistant Flag-Cdt2 (T464A, susceptible to FBXO11). We then monitored the effect on their half-lives by overexpressed myc- Mdm2. The results indicate that Mdm2 reduces half-life of both the mutants, suggesting that Mdm2-mediated destabilization of Cdt2 occurs independently of T464 phosphorylation (**Fig. 3E-F**).

### Mdm2 interacts with- and polyubiquitinates Cdt2

If Mdm2 destabilizes Cdt2 through polyubiquitination, we would expect the two proteins to physically interact. Immunoprecipitation of endogenous Mdm2 from lysates of MG132-treated cells co-precipitated endogenous Cdt2, indicating that the two proteins physically interact under physiological conditions (**Fig.4A**). Furthermore, ectopically expressed Flag-Cdt2 co-immunoprecipitated with myc-Mdm2, and reciprocally, ectopically expressed Myc-Mdm2 co-immunoprecipitated with Flag-Cdt2 in 293T cell lysates (**Fig. 4B-C**). Treatment with the proteasome inhibitor MG132 increased the overall levels of both proteins and enhanced the stability of their interaction (**Fig. 4B-C**).

**Fig. 4.**
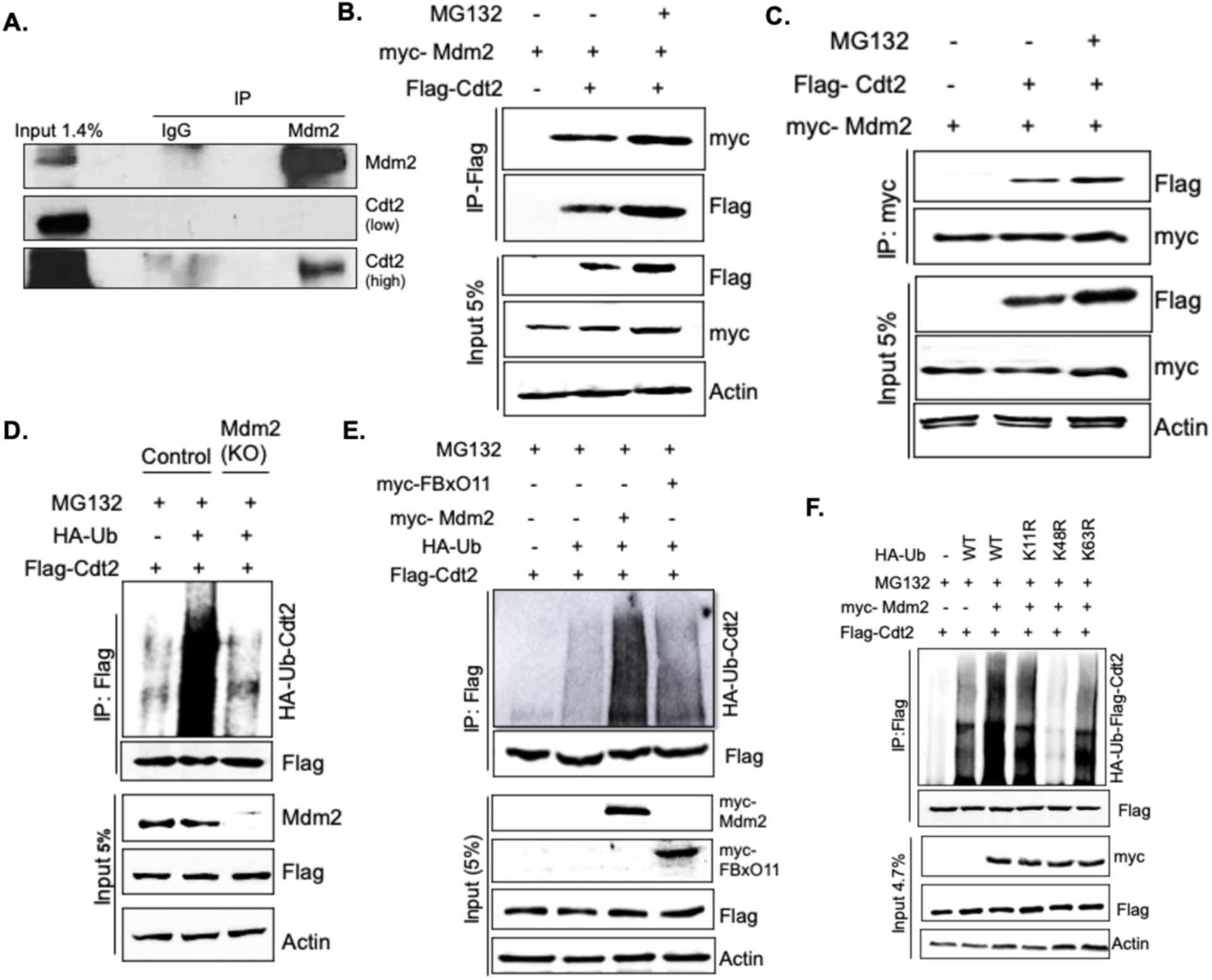
Mdm2 interacts with Cdt2 and promotes its ubiquitination via K48 linkage. (**A**) Coimmunoprecipitation of endogenous Mdm2 and Cdt2. Cell extracts from U2OS cells were immunoprecipitated (IP) with control IgG or Mdm2 antibodies, and the precipitates immunoblotted with the antibodies indicated to the right. (**B**) 293T cells were co-transfected with Flag-Cdt2 and myc-Mdm2 expressing plasmids as shown. Before harvesting cells were treated with DMSO or MGI32 (40 μM) for 6 h. The cell lysates were immunoprecipitated with anti-Flag antibodies. The Flag- immnoprecipitates were subjected to western blotting using antibodies indicated. (**C**) Coimmunoprecipitation of myc-Mdm2 and Flag Cdt2 as described in **B**. The extracts were immunoprecipitated with anti-myc antibodies and subjected to western blotting with antibodies indicated. (**D**) Mdm2 KO increases Cdt2 ubiquitination. Wild type or Mdm2 KO U2OS cells were treated with Flag-Cdt2 and HA-Ub plasmids. At 48 h after plasmid transfection, MG132 (40 μM) was added for 1 h before cells were harvested in denaturing ubiquitination buffer. The lysates were immunoprecipitated with anti-Flag antibody and probed first with anti-HA antibody to detect ubiquitinated forms of Cdt2 and then with anti-Flag antibodies to detect the immunoprecipitated Flag-Cdt2. (**E**) Overexpressing Mdm2 protein increases Cdt2 ubiquitination. 293T cells were transfected with plasmids as indicated. myc-FBxO11, a previously characterized E3 ligase for Cdt2 was used as a + control. The procedure was otherwise as described for panel D. (**F**) 293T cells were transfected with Flag-Cdt2, control or myc-Mdm2 along with wild type HA-ubiquitin plasmid or its different mutants as indicated. The ubiquitination assay for Flag-Cdt2 was performed as discussed for panel D.

To investigate whether Mdm2 promotes the polyubiquitination of Cdt2, we assessed the polyubiquitination levels of Flag-Cdt2 in wild-type (WT) and Mdm2 knockout (KO) cells. Under Mdm2 KO conditions, the polyubiquitination of Flag-Cdt2 was markedly reduced compared to wild-type cells (**Fig. 4D**), suggesting that Mdm2 is important for the polyubiquitination of Cdt2. Additionally, the polyubiquitination of Flag-Cdt2 was more robust when Mdm2 was overexpressed than when FBxO11 was overexpressed (**Fig. 4E**).

Furthermore, different types of polyubiquitin chains on substrates are formed by linking multiple ubiquitin molecules through one of their seven lysine residues. Analysis of polyubiquitin chain formation using various ubiquitin lysine-to-arginine mutants suggested that Mdm2 mediates K48-linked ubiquitination of Cdt2 (**Fig. 4F**). K48-linked polyubiquitin chains typically promote the ubiquitin-mediated degradation of substrate proteins.

Taken together, these results suggest that Mdm2 interacts with and polyubiquitinates Cdt2 via K48-linked chains.

### The degradation of Cdt2 by Mdm2 stabilizes CRL4^Cdt2^ substrates during G2/M

The results presented above suggest that Mdm2 may play a role in regulating cell cycle progression at the G2/M phase (Fig. 1B-D). The CRL4^Cdt2^ E3 ubiquitin ligase similarly regulates the cell cycle by targeting its substrates: p21, Set8, and Cdt1 for degradation during G1/S phase. These substrates typically reappear during the G2/M phase (Coleman et al. 2015). Upon discovering that Mdm2 promotes Cdt2 degradation, we hypothesized that Mdm2 may antagonize the E3 ligase activity of CRL4^Cdt2^ in order to stabilize its substrates during G2/M to facilitate proper cell cycle progression.

Analysis of whole-cell lysates revealed that Mdm2 knockout (KO) results in accumulation of Cdt2 and a concomitant reduction in the levels of CRL4^Cdt2^ substrates, including p21, Set8, and Cdt1 (**Fig.5A**). Conversely, ectopic expression of Mdm2 leads to reduced Cdt2 levels and a corresponding increase in these substrates (**Fig. 5B**). Notably, changes in p53 levels induced by either Mdm2 knockout or overexpression do not affect the levels of these substrates (**Fig. 5A-B**).

**Fig. 5.**
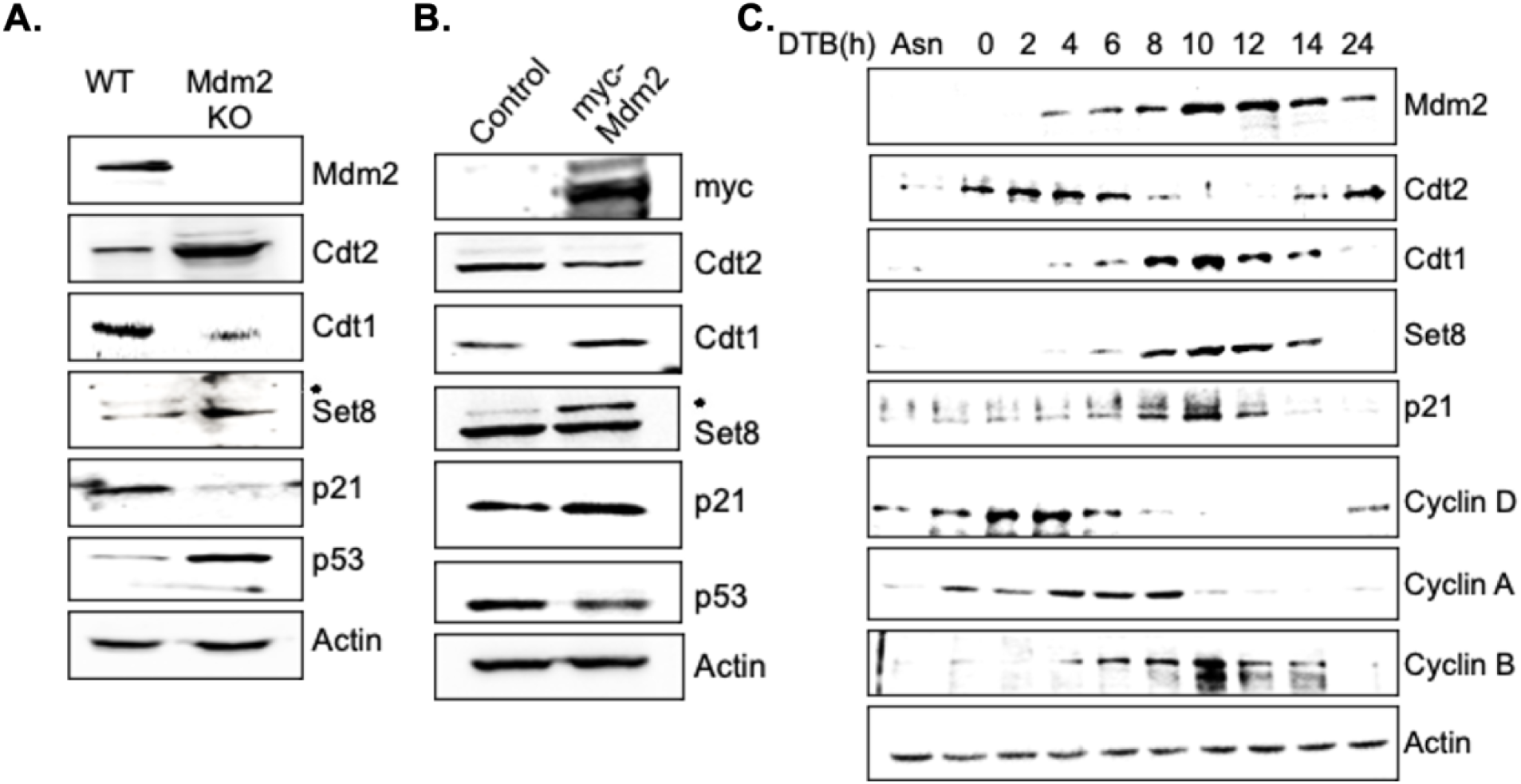
Mdm2 inactivates CRL4^Cdt2^ activity during G2/M. (A) Mdm2 KO increases Cdt2 and destabilises CRL4Cdt2 substrates. Western blot analysis of WT and Mdm2 KO U2OS cells with antibodies indicated. (B) Ectopic expression of myc-Mdm2 in U2OS cells destabilizes Cdt2 and as a consequence stabilizes Cdt2 substrates. The cell lysates are probed with antibodies indicated. (C) HeLa cells were synchronized by double thymidine block (DTB) and subsequent released. The cells were harvested at different time points followed by immunoblotting of lysates with antibodies specific for the indicated proteins. Asterisk indicates Set8 band.

To determine whether the re-emergence of CRL4^Cdt2^ substrates during G2/M correlates with the reciprocal dynamics of Mdm2 and Cdt2 expression, we synchronized HeLa cells at the G1/S boundary using a double thymidine block (DTB). Upon release from DTB, cells progressed through S phase at 4-6 hours, G2 at 8 hours, G2/M at 10 hours, and exited mitosis shortly thereafter, as determined by the temporal expression patterns of cyclin D, cyclin A, and cyclin B.

During G1/S and S phases, low Mdm2 expression coincides with elevated Cdt2 levels, resulting in the degradation of CRL4^Cdt2^ substrates. Conversely, as cells transition into G2, G2/M, and M phases, Mdm2 expression increases, leading to Cdt2 degradation and subsequent stabilization of key substrates such as Cdt1, Set8, and p21 (**Fig. 5C**). Together, these findings suggest that fluctuating Mdm2 levels orchestrate a biphasic regulation of CRL4^Cdt2^ substrate stability: promoting degradation during G1/S and S phases via Cdt2 activity when Mdm2 is low, and enabling substrate accumulation during G2/M through targeted Cdt2 degradation when Mdm2 levels peak. This regulatory mechanism highlights the critical role of Mdm2 in modulating CRL4^Cdt2^ activity and maintaining proper cell cycle progression.

### Mdm2-mediated neutralization of CRL4^Cdt2^ is critical for G2/M transition

As discussed above, the absence of Mdm2 impairs cell proliferation by causing cell cycle arrest at the G2/M phase (**Fig. 1B-C**), in a p53-independent manner. The marked upregulation of Cdt2 and the consequent downregulation of its substrates in Mdm2 knockout (KO) cells suggest that G2/M arrest and reduced proliferation may result from aberrant CRL4^Cdt2^ activity. Given that Mdm2 KO leads to destabilization of CRL4^Cdt2^ substrates, we investigated whether the observed G2/M arrest and reduced cell proliferation could be reversed by stabilizing these substrates. To this end, we reduced Cdt2 levels via shRNA-mediated knockdown in Mdm2 KO cells. As shown in **Fig. 6A**, silencing Cdt2 in Mdm2-deficient cells led to the stabilization of p21, Set8, and Cdt1. Moreover, Cdt2 knockdown partially rescued the G2/M arrest induced by Mdm2 loss (**Fig. 6B-C**). In line with these findings, the reduced proliferation observed upon Mdm2 KO was also partially restored by transient Cdt2 knockdown (**Fig 6D**). Collectively, these results demonstrate that Mdm2-dependent degradation of Cdt2 plays a crucial role in facilitating the G2/M transition.

**Fig. 6.**
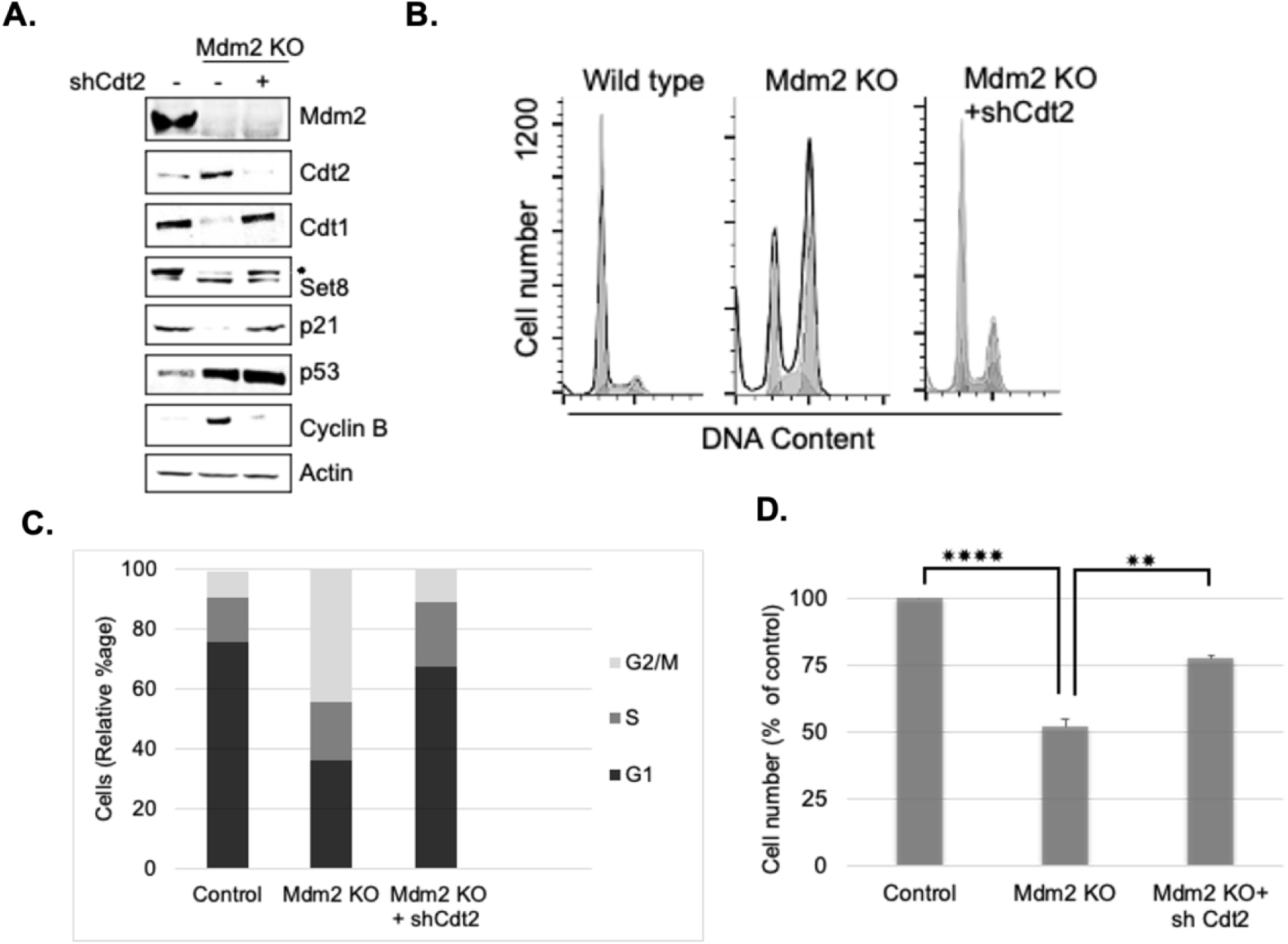
Mdm2 promotes G2/M transition and cell proliferation by targeting Cdt2 for degradation. **(A)** Silencing Cdt2 in Mdm2 KO U2OS cells rescues Cdt2 substrates. Lysates were prepared from Wild type, Mdm2 KO and Mdm2 KO+ shCdt2 cells and probed with indicated antibodies. (**B**) Fluorescence-activated cell sorting (FACS) profile of different cell cycle phases of Wild type, Mdm2 KO and Mdm2 KO+shCdt2 U2OS cells. (**C**) The relative percentage of cells used in Panel B are plotted. (**D**) MTT based cell viability assay indicating that co-depletion of Cdt2 rescues growth arrest in Mdm2 KO cells.

### p21 is the key effector of Mdm2-mediated G2/M transition and cell proliferation

Previous data demonstrated that key substrates of the CRL4^Cdt2^ complex: p21, Set8, and Cdt1, are destabilized in the absence of Mdm2 (**Fig. 6A**), and that transient depletion of Cdt2 in Mdm2 knockout (KO) cells alleviates the associated cell cycle defects, likely due to the stabilization of these substrates. To identify the specific substrate(s) responsible for the observed rescue in cell proliferation and G2/M progression, combinatorial knockdown experiments were performed in Mdm2 KO cells. Cdt2 was silenced in combination with individual knockdowns of Cdt1, Set8, or p21 using specific shRNAs (**Fig. 7A**). Co-depletion of either Cdt1 or Set8 did not affect the proliferative rescue mediated by Cdt2 knockdown. In contrast, co-silencing of p21 significantly impaired the proliferation rescue effect of Cdt2 depletion (**Fig. 7A-B**). Notably, a similar inhibitory effect was observed when all three substrates were simultaneously depleted along with Cdt2 (**Fig. 7A-B**).

**Fig. 7.**
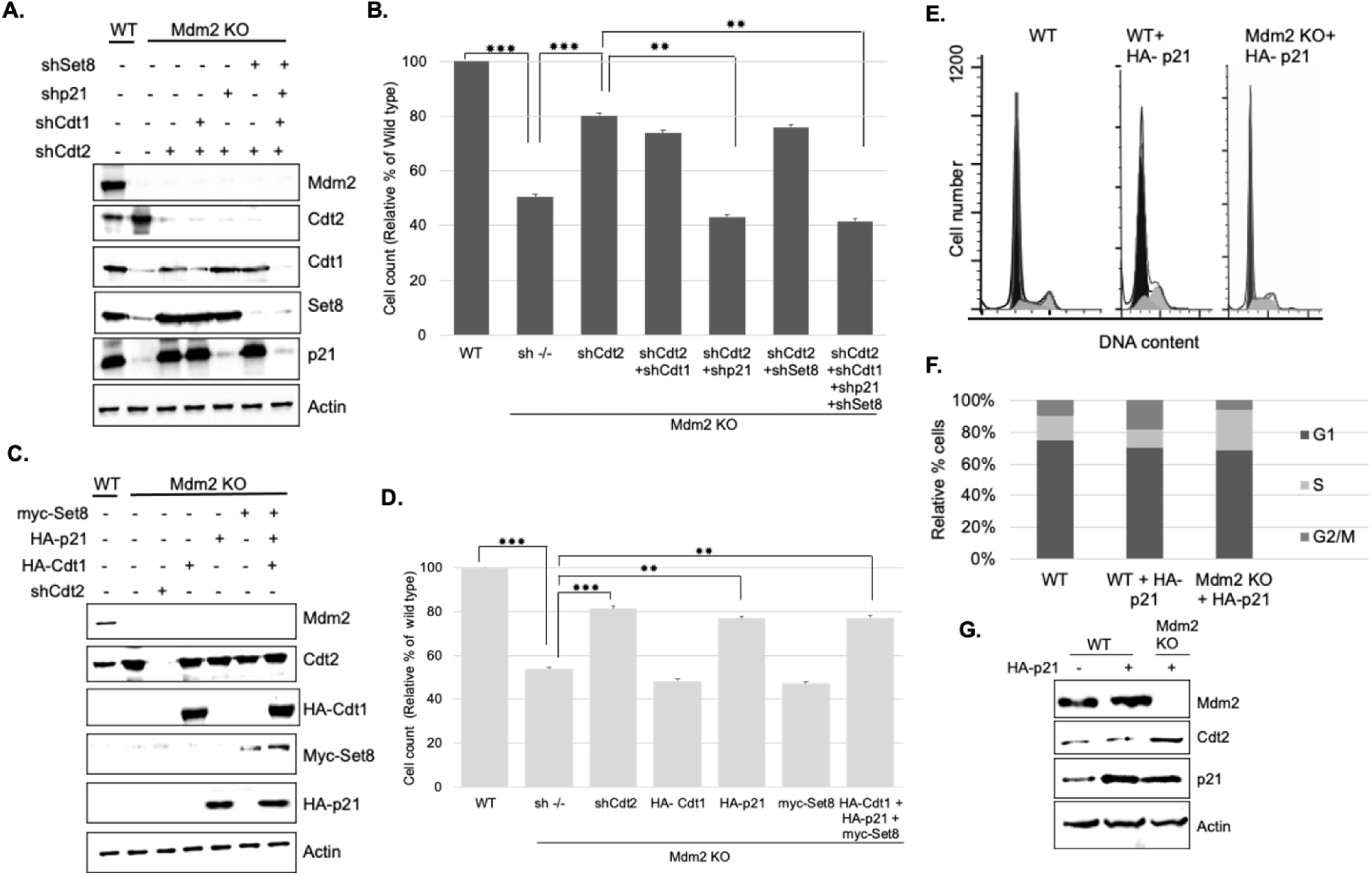
p21 is essential for Mdm2-dependent G2/M transition and cell proliferation. (**A**) Combinatorial knockdown of shCdt2 and its substrates (Cdt1, p21, Set8) in Mdm2 knockout (KO) U2OS cells. Mdm2 KO cells were transfected with shCdt2 alone or in combination with shCdt1, shp21, or shSet8, either individually or together, as indicated. After 60 hours of transfection, both wild-type and KO cells were harvested, and protein levels were analysed by western blotting using the indicated antibodies.(**B**) Cells grown in parallel to the experiment in panel A were used to determine cell viability using MTT assay. (**C**) Combinatorial overexpression of HA- Cdt1, HA-p21, myc-Set8 in Mdm2 knockout (KO) U2OS cells. Mdm2 KO cells were transfected with shCdt2 or plasmids overexpressing HA-Cdt1, HA-p21, or myc-Set8, either individually or together, as indicated. After 48 hours of transfection, both wild-type and KO cells were harvested, and protein levels were analysed by western blotting using the indicated antibodies. (**D**) Cells grown in parallel to the experiment in panel C were used to determine cell viability using MTT assay. (**E**) FACS analysis of different cell cycle phases of Wild type, wild type cells overexpressing HA-p21, and Mdm2 KO cells overexpressing HA-p21. (**F**) The relative percentage of cells used in Panel E are plotted. (**G**) Cell lysates from experiments performed in parallel to those whose results are shown in panels E, and F were probed with the indicated antibodies to monitor the p21 levels.

To further validate these findings, we ectopically expressed HA-Cdt1, myc-Set8, or HA-p21 in Mdm2-deficient cells and assessed their ability to restore cell proliferation. While Cdt2 knockdown effectively rescued the proliferation defect in Mdm2 KO cells, ectopic expression of Cdt1 or Set8 failed to confer any rescue effect. In contrast, reintroduction of HA-tagged p21 significantly restored the proliferative capacity of Mdm2-deficient cells (**Fig. 7C-D**).

Next, we examined whether overexpression of HA-p21 could overcome the G2/M arrest observed in Mdm2 KO cells, despite elevated Cdt2 levels. As expected, HA-p21 overexpression had no significant effect on cell cycle progression in normal cells (**Fig. 7E-G**). However, in Mdm2-deficient cells, HA-p21 overexpression successfully rescued the G2/M block (**Fig. 7E-G**), indicating that the stability of p21, regulated by Mdm2 through the CRL4^Cdt2^ pathway, is critical for G2/M transition.

Collectively, these results identify p21 as a key effector among CRL4^Cdt2^ substrates regulated by Mdm2. They highlight the essential role of p21 in promoting G2/M transition and cell proliferation in a p53-independent manner, emphasizing the importance of Mdm2-mediated p21 stabilization in maintaining proper cell cycle progression.

## Discussion

Mdm2, a critical oncoprotein, has been extensively studied for its role in regulating p53 (Haupt et al. 1997). However, since approximately 50% of cancers harbour mutations or deletions in p53 gene (Joerger and Fersht 2016), it becomes increasingly important to investigate p53-independent mechanisms by which Mdm2 may promote oncogenesis. This study positions Mdm2 as a key regulator of the G2/M phase transition and cellular proliferation independently of p53 (**Fig. IB-C**). Mdm2 carries out these functions by facilitating the re-accumulation of the cell cycle regulator p21 after its degradation in S-Phase by CRL4^Cdt2^ E3 ligase. We show that Mdm2- mediated regulation of p21 stability is a critical determinant of G2/M phase progression and cell proliferation (**Fig. ID, 7A-G**).

Mdm2 has previously been linked to cell cycle progression through regulation of Cyclin A levels (Frum et al. 2009) or paradoxically inhibition of the cell cycle regulator Cdc25A (Giono et al. 2017). However, those roles are either p53-dependent or occur in response to DNA damage. The genetic ablation of Mdm2 in U2OS cells with CRISPR-Cas9 technology allowed us to delineate the p53’s influence on Mdm2’s regulatory role in cell cycle under basal conditions (**Fig 1A-C**). Our observation that Mdm2 KO induces G2/M arrest is consistent with the idea that Mdm2 may have a 53- independent role in cell cycle progression by alternately regulating the expression of p73 and E2F family members (Klein et al. 2021a). Here, we determine that, Mdm2 regulates the CRL4^Cdt2^ E3 ubiquitin ligase complex during the G2/M phase by targeting Cdt2 subunit (and DDB1 subunit) for proteasomal degradation via K48-linked ubiquitination independent of p53 (**Fig. 2G**, **4F**).

Earlier studies have shown that Cdt2 is susceptible to ubiquitin-mediated degradation by SCF^FBXO11^ upon cell cycle exit or in response to stress signals (e.g., TGF-β signalling), which trigger its degradation. Phosphorylation of the Thr464 residue by CDKs (Cyclin B-Cdk1/Cyclin A-Cdk2) delays this process (Rossi et al. 2013)(Abbas et al. 2013). Additionally, 14-3-3 proteins bind to phospho-Thr464, shielding Cdt2 from FBXO11-mediated degradation (Dar et al. 2014). Strikingly, Mdm2 promotes proteasomal degradation of phosphomimetic ( T464D) and phosphorylation-defective (T464A) Cdt2 mutants, suggesting that Mdm2-mediated degradation occurs independently of T464 phospho-switch to facilitate the G2/M transition (**Fig. 3E-H**). This distinct degradation pathway for Cdt2 potentially involves alternate binding interfaces or post translational modifications that need explored investigation.

The oscillatory expression pattern of Mdm2 during the cell cycle further supports its role as a gatekeeper of CRL4^Cdt2^ activity. Low Mdm2 levels during S-phase coincide with Cdt2 accumulation and active degradation of CRL4^Cdt2^ substrates such as p21, Set8 and Cdt1. Conversely, elevated Mdm2 during G2/M (Gu et al. 2003) promotes Cdt2 degradation, allowing the accumulation of these substrates (**Fig. 5C**). Notably, the forced depletion of Cdt2 in Mdm2-deficient cells restores these substrates and partially rescues both G2/M arrest and impaired proliferation, suggesting the importance of downregulation of Cdt2 for G2/M phase progression (**Fig. 6 A-D**).

Among CRL4^Cdt2^ substrates, the accumulation of p21 plays a central role in the G2/M transition. By co-silencing Cdt2 along with individual substrates (Cdt1, p21, Set8) in Mdm2 knockout (KO) cells, we developed a system to identify which stabilized substrate is critical for G2/M progression and cell proliferation. This analysis identified p21,but not Set8 or Cdt1, as the key determinant of Mdm2-mediated cell cycle progression (**Fig.7A-B**). Furthermore, ectopic expression of p21 in the Mdm2 KO background further distinguishes p21 from Cdt1 and Set8 in its requirement for cell cycle progression and underscores the functional significance of the Mdm2-CRL4^Cdt2^- p21 axis in regulating the G2/M phase and maintaining cell viability (**Fig 7C-G**). These findings also highlight the unique role of p21 in promoting G2/M transition (**Fig. 7E-F**) and cell proliferation under homeostatic conditions, distinct from its canonical function as a p53 effector in DNA damage-induced checkpoints. Our results support a growing body of evidence pointing to non-traditional, pro-mitotic, tumor-promoting roles for p21, including its ability to inhibit apoptosis and support mitotic progression, particularly in p53-deficient or stressed cancer contexts (Kreis et al. 2015)(Kreis et al. 2019)(Gartel and Tyner 2002)(Abbas and Dutta 2009). Furthermore, the pro-survival role of p21 in non-small cell lung carcinoma within a p53-wild-type context supports our findings (Cutty et al. 2025).

Taken together, our data establish Mdm2 as a master regulator of the CRL4^Cdt2^ complex and its substrates, orchestrating a proteolytic switch that ensures proper progression through the G2/M phase. This function is mechanistically distinct from Mdm2’s well-characterized role in p53 regulation and reveals a broader role for Mdm2 in ubiquitin-mediated control of the cell cycle machinery. Given the frequent overexpression of Mdm2 in human cancers (Oliner et al. 1992)(Oliner et al. 2016), these findings suggest that Mdm2 may promote tumorigenesis not only by suppressing p53 but also by disrupting the finely tuned degradation dynamics of Cdt2 and its substrates.

Future studies should investigate the impact of this pathway in tumor models where Mdm2 is overexpressed, regardless of p53 status-a context in which the non-canonical Mdm2-CRL4^Cdt2^-p21 pathway may be particularly relevant. This also highlights the importance of pharmacologically targeting Mdm2, independent of p53 activation, as a potential strategy for treating malignancies (Klein et al. 2021b).

## Competing interest statement

The authors declare no competing interests.

## Acknowledgments

This work was supported by DST-SERB grant EMR/2016/005111 and DBT Ramalingaswami fellowship awarded to A.D. The authors thank Dr. Anindya Dutta, University of Alabama for advice. The authors acknowledge the FIST support from the ANRF, India to the Department of Biochemistry and DBT-Builder for infrastructural support to the University of Kashmir.

---

**Fig S1:**
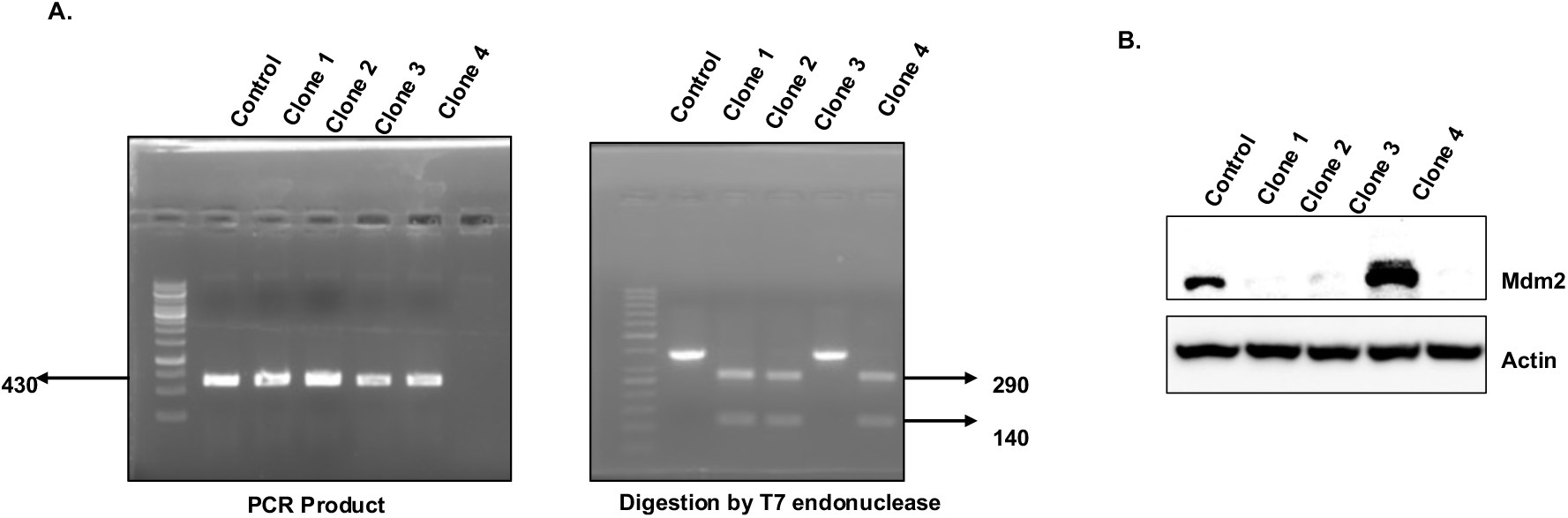
Mdm2 gene knockout using CRISPR-Cas9 . (A) PCR on genomic DNA of indicated clones. Digestion of PCR product from different clones with T7 endonuclease demonstrates clone 2, 3 and 4 have biallelic deletion of Mdm2 gene. (B) Verification of Mmd2 KO clones using western blotting with Mdm2 antibodies.

**Fig S2:**
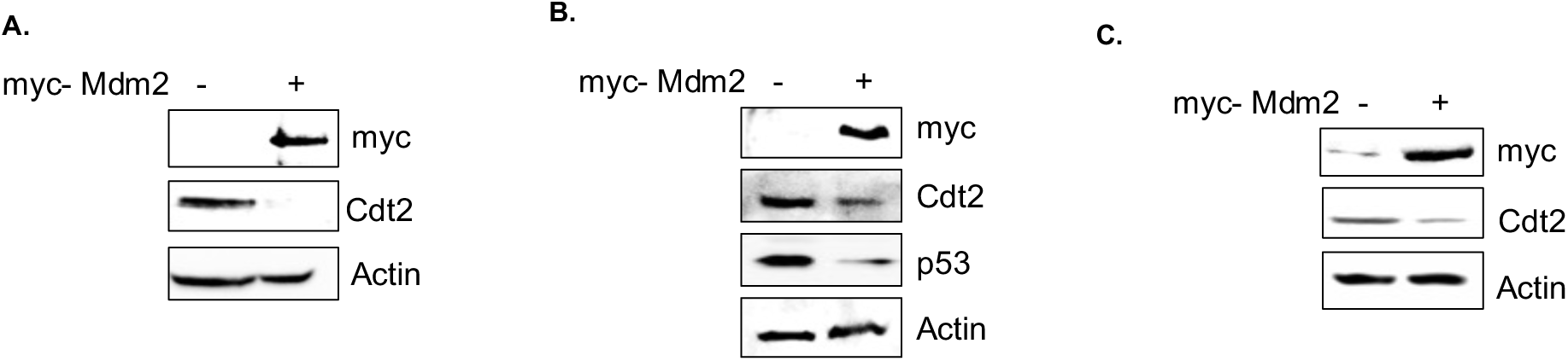
Ectopic myc-Mdm2 decreases endogenous Cdt2 levels in different cell lines. The results of western blot analysis of (A) 293T, (B) HCT116 and (C) HeLa cells, transfected with plasmid expressing control or myc-Mmd2 in are shown.

**Fig S3:**
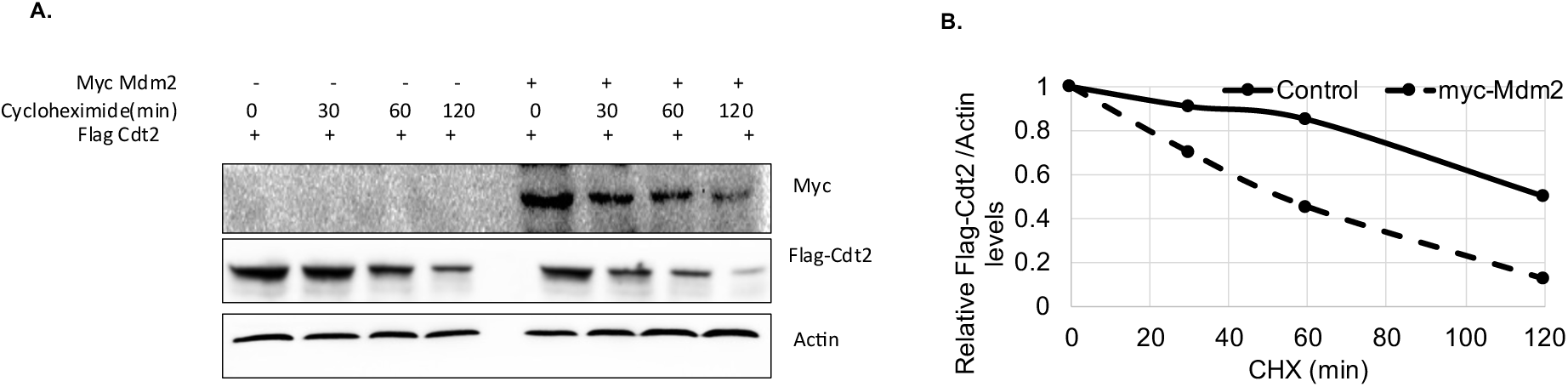
Mdm2 decreases the half-life of Cdt2. (**A**) The plasmid encoding Flag-Cdt2 was coexpressed in 293T cells with empty or myc-Mdm2 plasmids followed by treating cells with CHX for different time points. The cell lysates were probed with antibodies shown. (**B**) The Flag- signals in panel E were quantified, normalized to actin and plotted graphically.

**Table S1:**
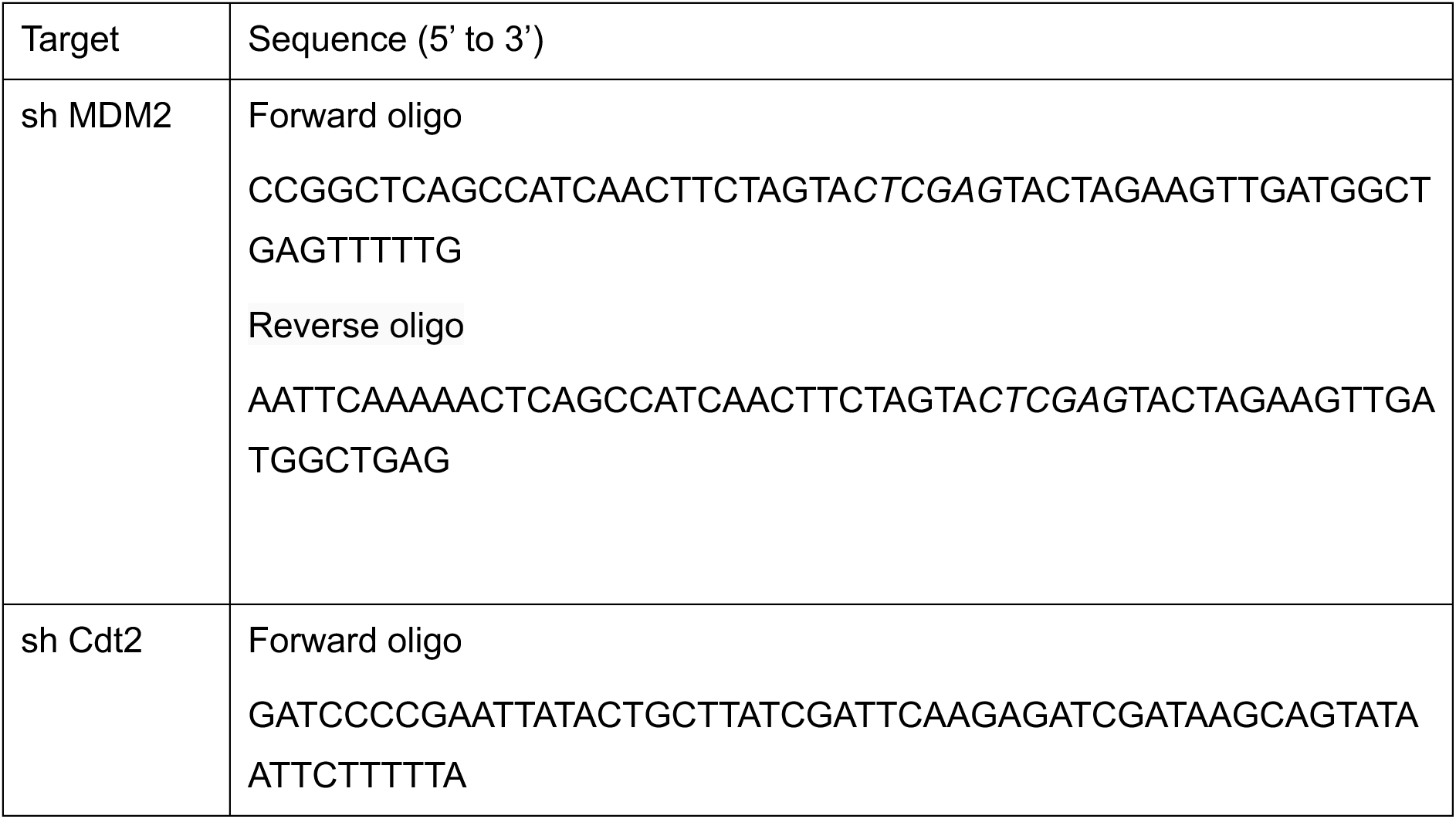
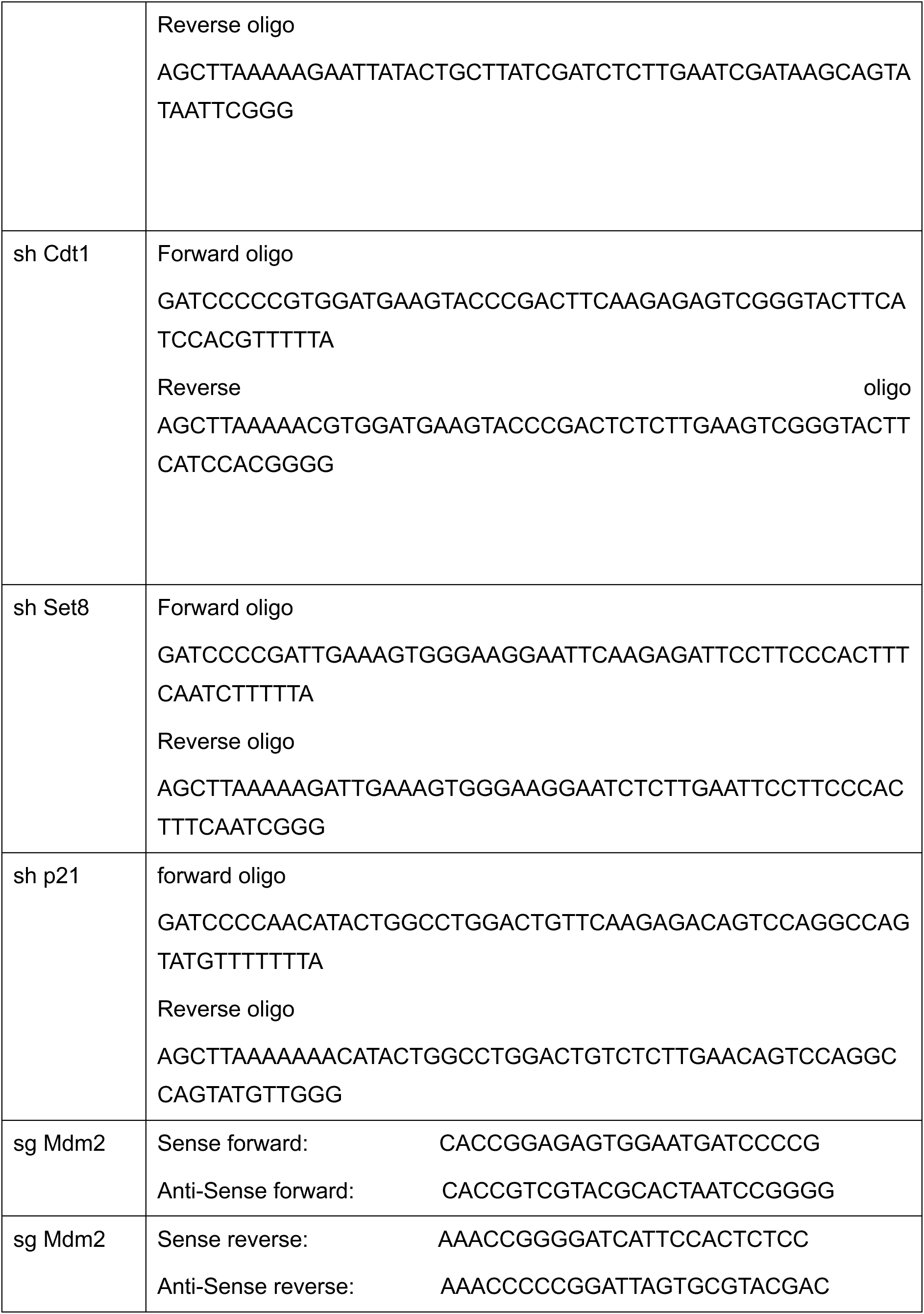
Sequences of Oligos used:

## Notes

### Competing Interest Statement

The authors have declared no competing interest.

### Summary of Updates

The order of Authors has been changed

