## Supplemental Data 1 for "Mdm2 restrains the activity of the CRL4^Cdt2^ E3 ubiquitin ligase to promote cell cycle progression through the G2/M phase"

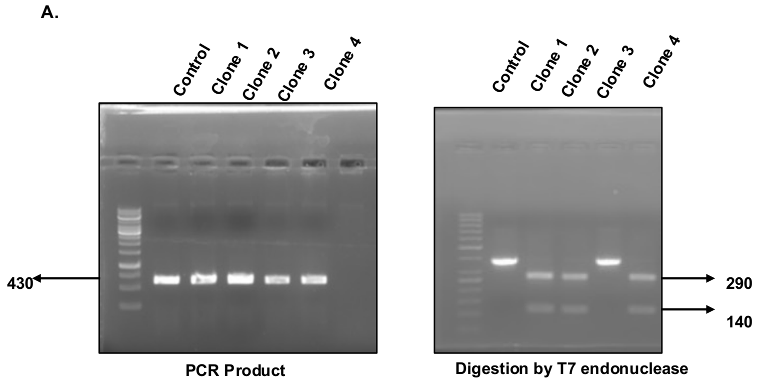


Fig S1: Mdm2 gene knockout using CRISPR-Cas9 . (A) PCR on genomic DNA of indicated clones. Digestion of PCR product from different clones with T7 endonuclease demonstrates clone 2, 3 and 4 have biallelic deletion of Mdm2 gene. (B) Verification of Mmd2 KO clones using western blotting with Mdm2 antibodies.

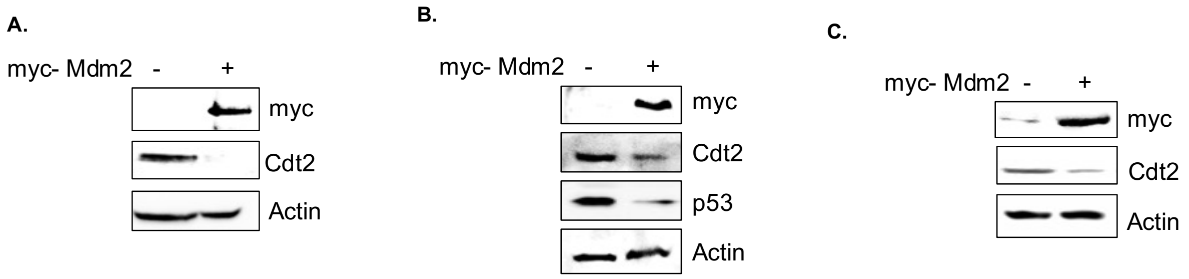

**Fig S2:** Ectopic myc-Mdm2 decreases endogenous Cdt2 levels in different cell lines . The results of western blot analysis of (A) 293T, (B) HCT116 and (C) HeLa cells, transfected with plasmid expressing control or myc-Mmd2 in  are shown.

Fig. S3

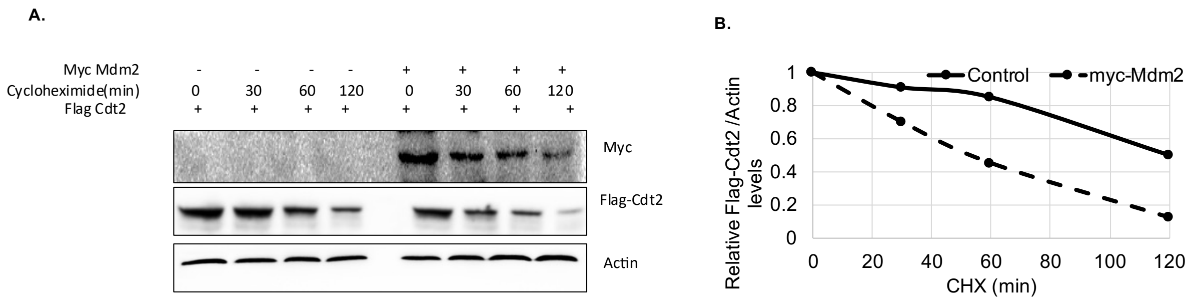


Fig S3: Mdm2 decreases the half-life of Cdt2. (A) The plasmid encoding  Flag-Cdt2 was coexpressed in 293T cells  with empty or myc-Mdm2 plasmids followed by treating cells with CHX for different time points. The cell lysates were probed with antibodies shown. (**B**) The Flag- signals in panel E were quantified, normalized to actin and plotted graphically.

Table S1: Sequences of Oligos used:

| Target | Sequence (5' to 3') |
|---|---|
| sh MDM2 | Forward oligo<br>CCGGCTCAGCCATCAACTTCTAGTA*CTCGAG*TACTAGAAGTTGATGGCTGAGTTTTTG<br>Reverse oligo<br>AATTCAAAAACTCAGCCATCAACTTCTAGTA*CTCGAG*TACTAGAAGTTGATGGCTGAG |
| sh Cdt2 | Forward oligo<br>GATCCCCGAATTATACTGCTTATCGATTCAAGAGATCGATAAGCAGTATAATTCTTTTTA |

| | |
|---|---|
| | Reverse oligo<br>AGCTTAAAAAGAATTATACTGCTTATCGATCTCTTGAATCGATAAGCAGTATAATTCGGG |
| sh Cdt1 | Forward oligo<br>GATCCCCCGTGGATGAAGTACCCGACTTCAAGAGAGTCGGGTACTTCATCCACGTTTTTA<br>Reverse oligo<br>AGCTTAAAAACGTGGATGAAGTACCCGACTCTCTTGAAGTCGGGTACTTCATCCACGGGG |
| sh Set8 | Forward oligo<br>GATCCCCGATTGAAAGTGGGAAGGAATTCAAGAGATTCCTTCCCACTTTCAATCTTTTTA<br>Reverse oligo<br>AGCTTAAAAAGATTGAAAGTGGGAAGGAATCTCTTGAATTCCTTCCCACTTTCAATCGGG |
| sh p21 | forward oligo<br>GATCCCCAACATACTGGCCTGGACTGTTCAAGAGACAGTCCAGGCCAGTATGTTTTTTTA<br>Reverse oligo<br>AGCTTAAAAAAACATACTGGCCTGGACTGTCTCTTGAACAGTCCAGGCCAGTATGTTGGG |
| sg Mdm2 | Sense forward: CACCGGAGAGTGGAATGATCCCCG<br>Anti-Sense forward: CACCGTCGTACGCACTAATCCGGGG |
| sg Mdm2 | Sense reverse: AAACCGGGGATCATTCCACTCTCC<br>Anti-Sense reverse: AAACCCCCGGATTAGTGCGTACGAC |
